# mTORC1 signaling is not essential for the maintenance of muscle mass and function in adult sedentary mice

**DOI:** 10.1101/738294

**Authors:** Alexander S. Ham, Kathrin Chojnowska, Lionel A. Tintignac, Shuo Lin, Alexander Schmidt, Daniel J. Ham, Michael Sinnreich, Markus A. Rüegg

**Affiliations:** Biozentrum, University of Basel, Basel, Switzerland; Department of Biomedicine, Pharmazentrum, University of Basel, Basel, Switzerland; Proteomics Core Facility, Biozentrum, University of Basel, Basel, Switzerland

**Keywords:** Raptor, Muscle atrophy, Protein translation, Fiber type, TOP mRNA

## Abstract

**Background:** The balance between protein synthesis and degradation (proteostasis) is a determining factor for muscle size and function. Signaling *via* the mammalian target of rapamycin complex 1 (mTORC1) regulates proteostasis in skeletal muscle by affecting protein synthesis and autophagosomal protein degradation. Indeed, genetic inactivation of mTORC1 in developing and growing muscle causes atrophy resulting in a lethal myopathy. However, systemic dampening of mTORC1 signaling by its allosteric inhibitor rapamycin is beneficial at the organismal level and increases lifespan. Whether the beneficial effect of rapamycin comes at the expense of muscle mass and function is yet to be established.

**Methods:** We conditionally ablated the gene coding for the mTORC1-essential component raptor in muscle fibers of adult mice (iRAmKO). We performed detailed phenotypic and biochemical analyses of iRAmKO mice and compared them with RAmKO mice, which lack raptor in developing muscle fibers. We also used polysome profiling and proteomics to assess protein translation and associated signaling in skeletal muscle of iRAmKO mice.

**Results:** Analysis at different time points reveal that, as in RAmKO mice, the proportion of oxidative fibers decreases, but slow-type fibers increase in iRAmKO mice. Nevertheless, no significant decrease in body and muscle mass, or muscle fiber area was detected up to 5 months post-raptor depletion. Similarly, *ex vivo* muscle force was not significantly reduced in iRAmKO mice. Despite stable muscle size and function, inducible raptor depletion significantly reduced the expression of key components of the translation machinery and overall translation rates.

**Conclusions:** Raptor depletion and hence complete inhibition of mTORC1 signaling in fully-grown muscle leads to metabolic and morphological changes without inducing muscle atrophy even after 5 months. Together, our data indicate that maintenance of muscle size does not require mTORC1 signaling, suggesting that rapamycin treatment is unlikely to negatively affect muscle mass and function.

## Introduction

Maintaining skeletal muscle is crucial for preserving the quality of life. Skeletal muscle atrophy, which occurs as a consequence of a range of conditions and diseases, such as immobilization, cancer cachexia or sarcopenia, the age-related loss of muscle mass and function, decreases independence and increases the risk of morbidity and mortality (Janssen et al., 2002; Lenk et al., 2010). Understanding the underlying mechanisms involved in maintaining skeletal muscle is a prerequisite for developing effective therapeutic strategies to prevent muscle atrophy. A protein complex postulated to be a key regulator in maintaining skeletal muscle mass is the mammalian (or mechanistic) target of rapamycin complex 1 (mTORC1) (Yoon, 2017). mTORC1 consists of the serine/threonine kinase mTOR, the regulatory-associated protein of mTOR (raptor), which is essential for the stability and function of the complex, and the mammalian lethal with Sec13 protein 8 (mLST8) (Laplante and Sabatini, 2012). Together, these proteins sense ATP, amino acid and insulin levels to promote or suppress protein translation (Laplante and Sabatini, 2012). mTORC1 promotes translation by releasing the eukaryotic translation initiation factor 4E (eIF4E) from its inhibitory binding partner eIF4E-binding protein 1 (4E-BP1) (Gingras et al., 2001), thereby allowing the formation of the eIF4F complex and recruitment of the pre-initiation complex to the 5’ end of mRNA (Ma and Blenis, 2009). Additionally, mTORC1 phosphorylates and activates ribosomal protein S6 kinase 1 (S6K1) that in turn phosphorylates S6, elongation factor 2 (eEF2) and initiation factor 4B (eIF4B) to promote protein synthesis (Ma and Blenis, 2009). Genetic or pharmacological inhibition of mTORC1 strongly decreases mRNA translation (Thoreen et al., 2012). In skeletal muscle, pharmacological inhibition or genetic ablation of crucial components of mTORC1 prevents overload-induced hypertrophy in rodents (Bentzinger et al., 2013; Bodine et al., 2001; You et al., 2019) and the mTORC1 inhibitor rapamycin blunts the amino acid-induced increase in protein synthesis in humans (Dickinson et al., 2011; Drummond et al., 2009). Furthermore, muscle fiber-specific knockout of raptor (Bentzinger et al., 2008) or mTOR (Risson et al., 2009) in developing mice reduces body and muscle mass, causes severe myopathic features and strongly reduces lifespan. Finally, ablation of raptor during myogenesis largely abrogates the formation of skeletal muscle (Rion et al., 2019). These results clearly demonstrate the importance of mTORC1 in skeletal muscle development and growth.

Based on these results, mTORC1 is also assumed to be a key regulator of muscle maintenance (Yoon, 2017). However, previous genetic studies that abrogated mTORC1 function were conducted during muscle growth (Bentzinger et al., 2013; Bodine et al., 2001; You et al., 2019) or used developmental knockout models (Bentzinger et al., 2008; Rion et al., 2019; Risson et al., 2009) where the phenotype may be the result from impairments during the rapid postnatal growth phase. Interestingly, long-term rapamycin treatment of adult mice does not appear to negatively affect whole-body muscle function (Neff et al., 2013; Zhang et al., 2014). The interpretation of these experiments is difficult, however, as long-term rapamycin treatment also affects mTORC2 function (Lamming et al., 2012) and because rapamycin only partially blocks mTORC1 (Thoreen et al., 2009). Therefore, the importance of mTORC1 in muscle maintenance is still unanswered. To analyze the role of mTORC1 specifically during adult muscle maintenance, we developed an inducible mouse model using the Tamoxifen/Mer-Cre-Mer system (McCarthy et al., 2012) where the gene for raptor (*Rptor*) can be ablated specifically in skeletal muscle during adulthood. By thoroughly examining the phenotype of these mice, from 10 days up to 5 months after raptor depletion, we demonstrate that, while these mice phenocopy some aspects of constitutive mTORC1 depletion in skeletal muscle, muscle mass and function are not affected for up to 5 months post-depletion. These results strongly indicate that muscle maintenance in sedentary mice is largely independent of mTORC1 activity and provide another example of the complex and context-dependent requirements of mTORC1 function.

## Methods

### Mice

All mice were kept under a 12-hour light to dark cycle in the animal facility of the Biozentrum (Basel, Switzerland) and received *ad libitum* access to water and standard laboratory chow. All experiments were approved by the veterinary commission of the Canton Basel-Stadt (Schlachthofstrasse 55, 4056 Basel, Switzerland).

Tamoxifen-inducible, muscle-specific raptor knockout mice (iRAmKOs) were generated by crossing three mouse lines. The mouse line with muscle-specific expression of Cre recombinase fused to two mutated estrogen receptors (*HSA-MerCreMer*) was generated and provided by Dr. K. Esser. (McCarthy et al., 2012) The modified *Rosa26* promoter with EGFP as a Cre reporter gene (*mR26-pCAG-EGFP^ki/ki^*) was generated and provided by Dr. M. Mueller (Tchorz et al., 2012). Homozygous *Rptor exon 6* floxed mice (*Rptor^fl/fl^*) were developed by our laboratory as previously described (Bentzinger et al., 2008). Experimental iRAmKO mice (*HSA-Mer-Cre-Mer^wt/ki^*; *Rptor^fl/fl^*; *mR26-pCAG-EGFP^ki/ki^*) and controls were generated by crossing *Rptor^fl/fl^*; *mR26-pCAG-EGFP^ki/ki^* with iRAmKO mice. *HSA-Mer-Cre-Mer^wt/ki^*-positive offspring were used as experimental mice (termed iRAmKOs), while Cre-negative mice were used as controls. To determine whether the mice were positive or negative for Cre we genotyped a toe-snip with forward (TGT GGC TGA TGA TCC GAA TA) and reverse primers (GCT TGC ATG ATC TCC GGT AT). All iRAmKOs and control mice were male.

### Tamoxifen administration

Tamoxifen powder (Sigma-Aldrich, St. Louis, MO, USA) was dissolved in corn oil (Sigma-Aldrich) at a concentration of 20 mg/ml. Tamoxifen was administrated *via* intraperitoneal injections of 100 µl when the mice were between 12 and 16 weeks old. The first administration regime comprised of five injections on five consecutive days (Supporting Information, Figure S1A). The day after the fifth injection was defined as day 1. Tamoxifen was administered again on day 3 and 4. For prolonged knockouts, two additional injections were administered on consecutive days every 28 days to prevent potential reintroduction of raptor into fibers by non-targeted muscle stem cells (MuSCs).

### Sample preparation and Western blot analysis

Muscles were powdered on a metal block cooled with liquid nitrogen. The tissue was lyzed in cold RIPA buffer (50 mM Tris-Base, 150 mM NaCl, 0.5% sodium deoxycholate, 0.1% SDS, 1% NP-40, all at pH 8) supplemented with a phosphatase (Roche, Basel, Switzerland) and protease inhibitor cocktail (Roche). The samples were incubated on a rotating wheel for 2 h at 4°C, sonicated twice for 8 s and centrifuged at 15,700 rcf for 30 minutes at 4°C. The protein concentration was measured on the cleared lysates with Pierce^TM^ BCA Protein Assay Kit (Thermo Fisher Scientific, Waltham, MA, USA). After diluting the lysates in RIPA and Laemmli buffer, equal protein amounts were loaded onto SDS gels (NuPage 4-12% Bis-Tris). After electrophoretic separation, proteins were transferred onto a nitrocellulose membrane on ice at 100 V for 1 h and stained with Ponceau to cut at the appropriate heights. The membrane slices were blocked with 4% (w/v) BSA in TBS-T solution (0.137 M NaCl, 2.68 mM KCl, 0.025 M-Tris-HCl pH 7.4, 0.1% Tween-20) for 45 min at RT. The primary antibodies were incubated over night at 4°C. After a washing step with TBS-T, the secondary antibodies were added for 1 h at RT. Western blot detection was done using the KLP LumiGlo Chemiluminescence Substrate Kit (Seracare, Milford, MA, USA) and imaged in a Fusion Fx machine (Vilber, Collégien, France). α-actinin was used as a housekeeping gene as well as for normalization.

### Antibodies used in Western blot

The following primary antibodies from Cell Signalling Technology (Danvers, MA, USA) were used: 4E-BP1 (#9452), P-4E-BP1^S65^ (#9451), AKT (#9272), P-AKT^S473^ (#9271), P-AKT^T308^ (#9275), eIF4E (#2067), mTOR (#2972), P-mTOR^S2448^ (#2971), raptor (#2280), S6 (#2217), S6^S240/244^ (#5364). The primary antibody for α-actinin (#A7732) was purchased from Abcam (Cambridge, England). The primary antibody for EGFP (11814460001) was from Roche Applied Science (Penzberg, Germany). All Cell Signalling Technology antibodies as well as EGFP were diluted 1:1,000, α-actinin was diluted 1:5,000. The secondary antibodies goat anti mouse (#115-035-003) and goat anti-rabbit (#111-035-003) were purchased from Jackson ImmunoResearch Europe (Ely, Cambridgeshire, England) and were diluted 1:10,000. All antibodies were diluted in 4% BSA in TBS-T.

### Immunohistochemistry

The muscles were frozen in 2-methylbutan directly after removal. All cross-sections were prepared on a cryostat at a thickness of 10 µm and stored at -20°C. For fiber type staining the sections were first rehydrated with PBS and then blocked for 30 min with blocking solution (0.4% Triton X-100, 10% goat serum [Biological industries, Beit Haemek, Israel] in PBS) followed by a washing step with PBS. The primary antibodies against MyHC 2b (1:100; #BF-F3; DSHB, Iowa, USA), MyHC 2a (1:200; #SC-71; DSHB), MyHC 1 (1:50; BA-D5; DSHB) and laminin (1:160; #ab11575; Abcam) were diluted in PBS containing 10% goat serum (Biological industries) and added for 1 h at RT. After a washing step with PBS, the secondary antibodies were added containing AF488 Gt anti-Ms IgM (1:100; Invitrogen, Carlsbad, CA, USA), AF568 Gt anti-Ms IgG1 (1:100; Invitrogen), DL405 Gt anti-Ms IgG2b (1:50; Jackson), AF647 Donkey anti-Rabbit IgG (1:200; Jackson) diluted in PBS containing 10% goat serum. The slides were again washed with PBS and then mounted with Vectashield Hardset (H-1600; Vector Labs, Burlingame, CA, USA). The entire section was imaged on a Zeiss (Oberkochen, Germany) Axio Scan.Z1 Slide Scanner.

### Real-time qPCR

Total RNA was isolated from the *soleus* using the Qiagen (Hilden, Germany) RNeasy Mini Kit. 500 ng of RNA was then used to generate first strand complimentary DNA (cDNA) using the iScript cDNA Synthesis Kit (Bio-Rad, Hercules, CA, USA). The cDNA was diluted 1:10 before usage and amplified on a LightCycler 480 (Roche Diagnostics, Basel, Switzerland) using the Applied Biosystems SYBR Green Master Mix (Roche) on a 384-well plate. The data was analyzed and quantified using the delta delta Ct method (Livak and Schmittgen, 2001). β-actin was used as a housekeeping gene as well as for normalization. The following primers were used: *Actb* fw: CAG CTT CTT TGC AGC TCC TT and bw: GCA GCG ATA TCG TCA TCC A; *Cox5b* fw: CTT CAG GCA CCA AGG AAG AC and bw: TTC ACA GAT GCA GCC CAC TA; *Cycs* fw: AAA TCT CCA CGG TCT GTT CG and bw: TAT CCT CTC CCC AGG TGA TG; *Fbxo32* fw: CTC TGT ACC ATG CCG TTC CT and bw: GGC TGC TGA ACA GAT TCT CC; *Mb* fw: ATC CAG CCT CTA GCC CAA TC and bw: GAG CAT CTG CTC CAA AGT CC; *Ppargc1α* fw: TGA TGT GAA TGA CTT GGA TAC AGA CA and bw: GCT CAT TGT TGT ACT GGT TGG ATA TG; *Trim63* fw: ACC TGC TGG TGG AAA ACA and bw: AGG AGC AAG TAG GCA CCT CA.

### Muscle force measurements

The *ex vivo* force measurement of the *extensor digitorum longus* (EDL) and the *soleus* was done as described previously (Reinhard et al., 2017). Briefly, muscles were carefully excised and mounted on the 1200A Isolated Muscle System (Aurora Scientific, Aurora, ON, Canada) in an organ bath containing 60 mL of Ringer solution (137 mM NaCl, 24 mM NaHCO_3_, 11 mM Glucose, 5 mM KCl, 2 mM CaCl_2_, 1 mM MgSO_4_, 1 mM NaH_2_PO_4_) that was gassed with 95% O_2_-5% CO_2_ at 30°C. Specific forces are derived by normalizing to the cross-sectional area as described (Brooks and Faulkner, 1988). The fatigue of EDL and *soleus* muscles was assessed by 6 min stimulation at 200 Hz and 120 Hz, respectively.

### m7GTP pulldown

The *gastrocnemius* was powdered on liquid nitrogen and lysed for 1 h at 4°C in lysis buffer (20 mM Hepes (pH 7.4), 50 mM KCl, 0.2 mM EDTA, 25 mM β-glycerophosphate, 0.5 mM sodium-orthovanadate, 1 mM dithiothreitol, 0.5 % Triton X-100, and 50 mM NaF) supplemented with a protease inhibitor cocktail tablet (Roche). After additional cell disruption with a potter homogenizer, the lysate was centrifuged for 10 min at 9,800 rcf at 4°C. The protein concentration was measured using the Pierce^TM^ BCA Protein Assay Kit (Thermo Fisher Scientific). A total of 400 µg was used for the pulldown. For pre-clearance 15 μl of blank Agarose beads (Jena Bioscience, Thüringen, Germany) were added for 30 min at 4°C. The samples were then centrifuged for 10 min at 9,800 rcf at 4°C. Next, twenty μl of a 50% slurry of m7GTP-Agarose beads (Jena Bioscience) was added to the supernatant overnight at 4°C. The beads were then washed three times using 200 μl of the lysis buffer for 10 min at 9,800 rcf. To finish, the samples were denatured in Laemmli buffer at 95°C before loading them for Western blot analysis.

### Fiber area and fiber size distribution

The fiber size distribution and the fiber-type composition was evaluated using the fiber-type staining described under Immunohistochemistry. Unstained fibers we defined as type 2X fibers. The entire muscle section was analyzed using a script on imageJ and Jupyter (Python). The variance coefficient was calculated using the minimal fiber Feret’s diameter as previously described (Briguet et al., 2004).

### Mass Spectrometry

Approximately 5 ug of *gastrocnemius* muscle tissue was collected, dissolved in 100 µl lysis buffer (1% sodium deoxycholate, 0.1M ammoniumbicarbonate), reduced with 5 mM TCEP for 10 min at 95°C and alkylated with 10 mM chloroacetamide for 30 min at 37°C. Samples were digested with trypsin (Promega, Fitchburg, WI, USA) at 37°C overnight (protein to trypsin ratio: 50:1) and desalted using iST cartridges according to the manufacturer’s instructions (Phoenix, PreOmics GmbH, Planegg, Germany).

One µg of peptides of each sample were subjected to LC–MS analysis using a dual pressure LTQ-Orbitrap Elite mass spectrometer connected to an electrospray ion source (both Thermo Fisher Scientific) as recently specified(Ahrne et al., 2016) and a custom-made column heater set to 60°C. Peptide separation was carried out on an EASY nLC-1000 system (Thermo Fisher Scientific) equipped with a RP-HPLC column (75μm × 30cm) packed in-house with C18 resin (ReproSil-Pur C18–AQ, 1.9 μm resin; Dr. Maisch GmbH, Ammerbuch-Entringen, Germany). Separation was achieved by a linear gradient from 95% solvent A (0.1% formic acid, 99.9% water) and 5% solvent B (80% acetonitrile, 0.1% formic acid, 19.9% water) to 28% solvent B over 75 min to 40% solvent B over 15 min to 95% solvent B over 2 min and 95% solvent B over 18 min at a flow rate of 0.2 μl/min.

The data acquisition mode was set to obtain one high resolution MS scan in the FT part of the mass spectrometer at a resolution of 240,000 full width at half maximum (at 400 m/z, MS1) followed by MS/MS (MS2) scans in the linear ion trap of the 20 most intense MS signals. The charged state screening modus was enabled to exclude unassigned and singly charged ions and the dynamic exclusion duration was set to 30s. The ion accumulation time was set to 300 ms (MS1) and 50 ms (MS2). MS1 and MS2 scans were acquired at a target setting of 1E6 ions and 10,000 ions, respectively. The collision energy was set to 35%, and one microscan was acquired for each spectrum.

To determine changes in protein expressions across samples, a MS1 based label-free quantification was carried out. Therefore, the generated raw files were imported into the Progenesis QI software (Nonlinear Dynamics, Version 2.0) and analyzed using the default parameter settings. MS/MS-data were exported directly from Progenesis QI in mgf format and searched against a decoy database of the forward and reverse sequences of the predicted proteome from *mus musculus* (Uniprot, download date: 2017/04/18, total of 34,490 entries) using MASCOT (version 2.4.1). The search criteria were set as following: full tryptic specificity was required (cleavage after lysine or arginine residues); 3 missed cleavages were allowed; carbamidomethylation (C) was set as fixed modification; oxidation (M) as variable modification. The mass tolerance was set to 10 ppm for precursor ions and 0.6 Da for fragment ions. Results from the database search were imported into Progenesis QI and the final peptide measurement list containing the peak areas of all identified peptides, respectively, was exported. This list was further processed and statically analyzed using our in-house developed SafeQuant R script (SafeQuant, https://github.com/eahrne/SafeQuant). The peptide and protein false discovery rate (FDR) was set to 1% using the number of reverse hits in the dataset.

The result details of the proteomics experiments carried out including identification scores, number of peptides quantified, normalized (by sum of all peak intensities) peak intensities, log2 ratios, coefficients of variations and p-values for each quantified protein and sample is displayed in Supporting Information, MS data.

### Polysome profiling

The *gastrocnemius* muscles were pulverized under cryogenic conditions in a cryo-freezer grinder (SpexSamplePrep, 10 cps 3x2min) in 2.5ml of lysis buffer (20 mM Tris-HCl, pH=7, 100 mM NaCl, 50 mM NH_4_Cl, 10 mM MgCl_2_, 1% Triton X-100 and freshly added 100 µg/ml cycloheximide, 1 mM dithiothreitol supplemented with a protease inhibitor cocktail tablet (Roche), and 200U of SUPERase*In (Invitrogen). After 2 h incubation at 4°C on a rotating wheel, lysates were disrupted by 5 up-and-down passages through a 19-gauge syringe and clarified by centrifugation for 10 min at 12,000 rcf at 4°C. Polysome profiling was further carried out as previously described by Ingolia and colleagues (Ingolia et al., 2013). Briefly, equal loading was assured by measuring OD260 for each sample. Lysates were then layered on-top of a pre-cooled 50-10% sucrose gradient (dissolved in 8.3 mM Tris-HCl, pH=7.5, 8.3 mM NH_4_Cl, 2 mM MgCl_2_ freshly supplemented with 0.083 mM dithiothreitol, 100 µg/ml cycloheximide and 200U of SUPERase*In). Gradient was formed using a Gradient Master instrument (Biocomp) according to the manufacturer’s instruction. Samples were centrifuged at 35,000 rpm for 3 h at 4°C using a Beckman SW41Ti rotor. Profiles of the different samples were obtained at a speed of 0.5 ml per min and continuous measurement of the absorbance at 254 nm.

## Results

### Generation of an inducible, muscle fiber-specific raptor knockout mouse

To investigate the role of mTORC1 signaling in adult muscle, we crossed *Rptor*^fl/fl^ mice (Bentzinger et al., 2008) with mice expressing a modified Cre-recombinase (Mer-Cre-Mer) under the control of the human skeletal actin (HSA) promoter (McCarthy et al., 2012). We name these mice <u>i</u>nducible <u>ra</u>ptor <u>m</u>uscle-specific <u>k</u>nock<u>o</u>ut (iRAmKO) mice. *Rptor* deletion was induced in three to four-month-old mice to assure that muscles had reached adult size. Mice were analyzed 10 days, 21 days, 3 months and 5 months after the induction of *Rptor* deletion. To monitor successful recombination, iRAmKO mice also carried an EGFP-reporter (Tchorz et al., 2012). Daily injections of tamoxifen (TAM) into iRAmKO mice for five consecutive days (Supporting Information, Figure S1A) caused translocation of Mer-Cre-Mer into the myonuclei resulting in recombination of the floxed alleles. The day after the fifth injection was defined as day 1 of the knockout (Supporting Information, Figure S1A). Ten days after TAM injection, all muscle fibers of the *tibialis anterior* (TA) were EGFP positive, indicating efficient recombination (Figure 1A and 1B; Supporting Information, Figure S1B). To assure complete loss of raptor protein, we also performed a detailed biochemical analysis of the mTOR pathway. Raptor protein levels were ∼30% of control after 10 days and ∼11% after 21 days and this low level of raptor remained constant after 3 months (Figure 1C and 1D). This residual raptor expression likely originates from non-targeted cells residing in skeletal muscle as has also been observed in constitutive, muscle fiber-specific RAmKO mice (Bentzinger et al., 2008). We also examined raptor protein levels in the heart to test tissue specificity. Despite a weak GFP signal, raptor levels were unchanged in the heart (Supporting Information, Figure S1B). In line with previous reports (You et al., 2019), loss of raptor protein significantly reduced total levels of mTOR as well as the phosphorylated (Serine2448) levels of mTOR protein (Figure 1C and 1D). Phosphorylation of the mTORC1 target 4E-BP1 was not altered at 10 days post-injection but was significantly lower 21 days and 3 months after TAM injection (Figure 1C and 1D). Levels of p-S6 showed a similar pattern as p-4E-BP1 but they did not reach statistical significance (Figure 1C and 1D). The activity of protein kinase B (PKB/AKT) is indirectly controlled *via* an inhibitory feedback loop involving the mTORC1 target S6K1 (Rozengurt et al., 2014). Accordingly, as previously observed in response to mTORC1 inhibition (Sarbassov et al., 2005), iRAmKO mice display higher AKT phosphorylation at 21 days and 3 months post-injection at both, the PDK1-dependent (p-AKT^T308^) and mTORC2-dependent site (p-AKT^S473^) (Figure 1C and 1D). In conclusion, these experiments show that TAM efficiently depletes raptor in iRAmKO mice leading to abrogation of mTORC1 signaling after 21 days.

**Figure 1.**
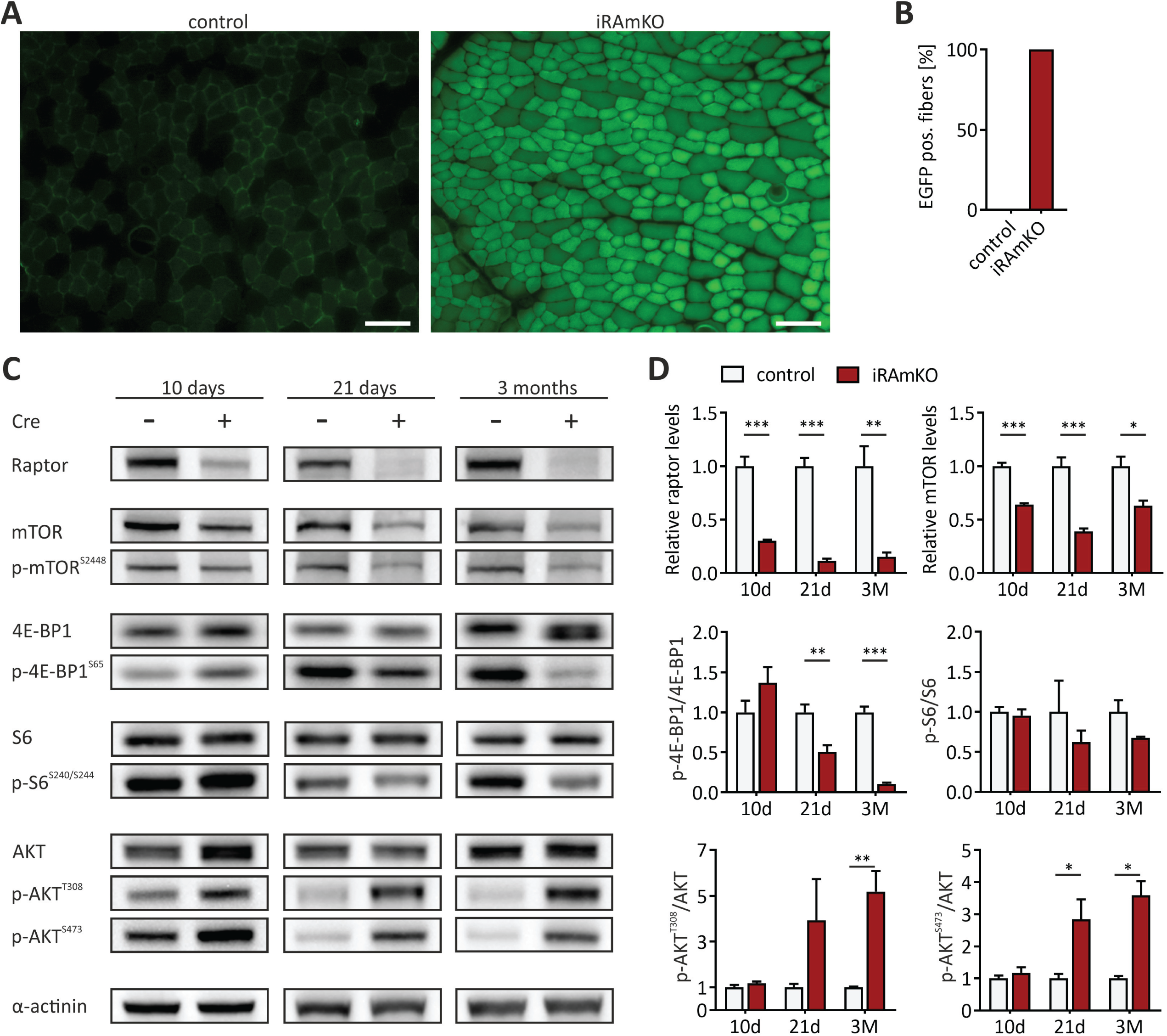
Generation of an inducible, muscle fiber-specific raptor knockout mouse. (A) TA muscle cross-sections of iRAmKOs 10 days post TAM treatment were fixed with PFA and imaged for GFP. Size bar = 100 µm. (B) Quantification of GFP positive fibers. (C) Western blot analysis of *tibialis anterior* (TA) lysates from three different time points after TAM injection. Equal amounts of protein were loaded in each lane and α-actinin was used as a loading control. Proteins detected in the different immunblots are indicated at the left. (D) Quantifications of the immunoblots shown in C. All values were normalized to α-actinin and mean values detected in Cre-negative (control) samples were set to 1. Quantifications for the indicated proteins were done 10 days (10d), 21 days (21d) or 3 months (3M) after TAM injection. 10d and 21d iRAmKOs n = 4, 3 month-iRAmKOs n ≥ 3 (Cre+ n = 3, Cre- n = 4). Values represent the mean ± SEM. Significance was assessed using two-tailed unpaired student’s *t*-test: *p < 0.05, **p < 0.01, ***p < 0.001.

### Prolonged depletion of raptor does not cause an overt phenotype

To analyze the long-term consequence of raptor depletion in adult muscle, we treated 3-month-old mice with TAM and analyzed the mice up to 5 months later. This time point was chosen as developmental, muscle-specific raptor knockout (RAmKO) mice show a severe myopathy at the age of 5 months and eventually succumb at the age of 5-6 months (Bentzinger et al., 2008). Contrary to what we expected, 5-month iRAmKO mice were barely distinguishable from their control littermates (Figure 2A). For example, 5-month iRAmKO mice did not display kyphosis, a sign of muscle weakness. Body mass of mutant mice was not different from littermate controls during the entire period (Figure 2B). Similarly, we could not detect any difference in relative fat mass after 3 months and only a slight, but significant, increase of fat mass after 5 months (Figure 2C). Lean mass was neither significantly changed at 3 nor at 5 months (Figure 2C). Weighing individual muscles showed that *extensor digitorum longus* (EDL), *soleus*, TA and *gastrocnemius* were not different to the control muscles with the exception of *soleus* muscle in 5-month iRAmKO mice, which was significantly heavier than the control (Figure 2D). Thus, removal of raptor in fully-grown muscle for up to 5 months does not promote loss of muscle or body mass.

**Figure 2.**
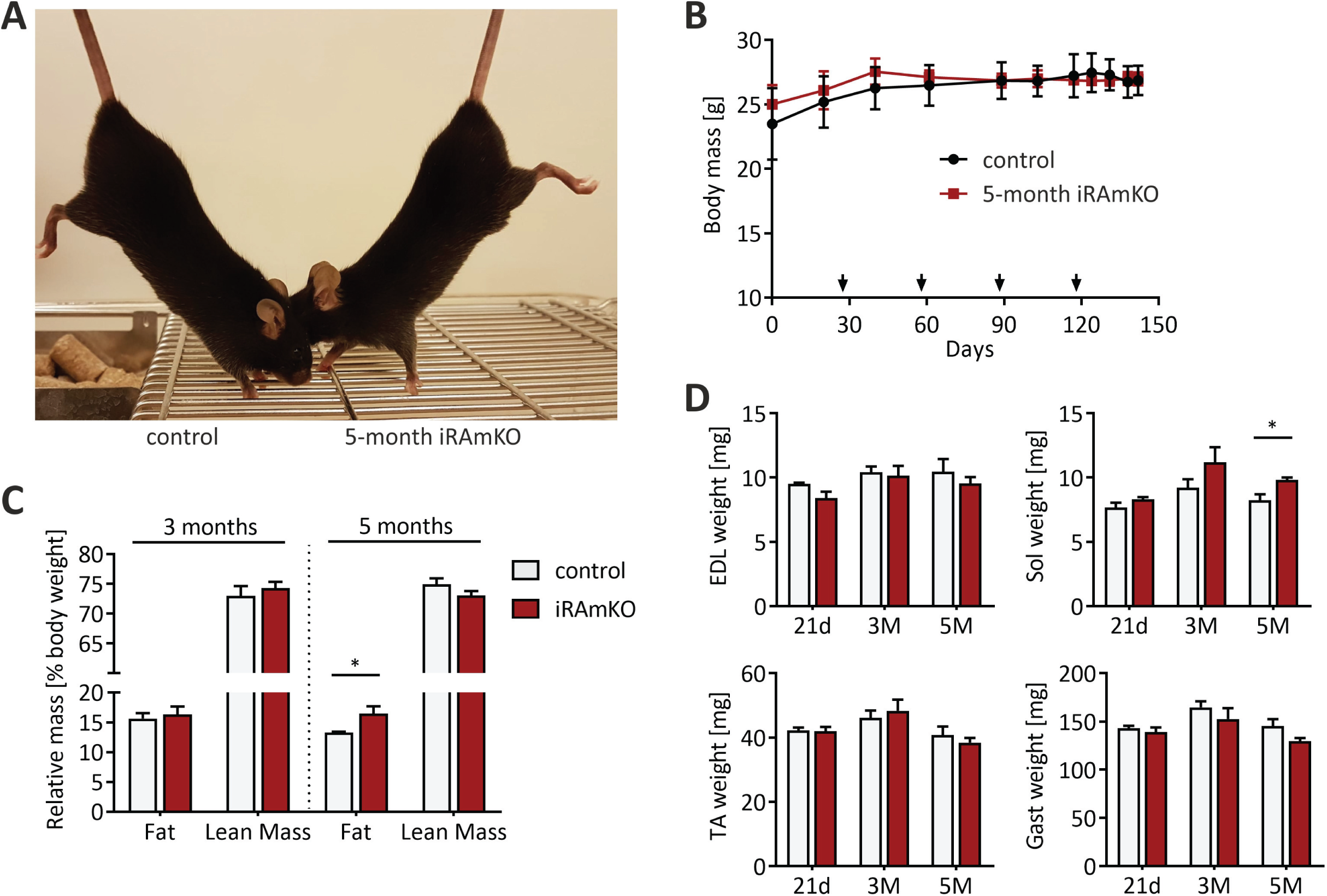
Prolonged depletion of raptor does not cause an overt phenotype. (A) Photograph of 8-month-old iRAmKO and control mouse that were 5-month depleted for raptor. (B) Weight curve of 5-month iRAmKOs and controls after TAM injection. The arrows indicate the monthly repeated TAM injections. At no time point was there a significant difference in the body mass. (C) Relative fat and lean mass measured via MRI normalized to body mass. 3-month iRAmKOs present no changes while 5-month iRAmKOs reveal an increase in fat mass. (D) The weight of the four hind limb muscles TA, *extensor digitorum longus* (EDL), *soleus* (Sol) and the *gastrocnemius* (Gast) were measured during dissection in 21-day, 3-months and 5-month iRAmKOs. 21-day and 5-month iRAmKOs n = 4, 3-month iRAmKOs n ≥ 3. Values represent the mean ± SEM. Significance was assessed using two-tailed unpaired student’s *t*-test: *p < 0.05, **p < 0.01, ***p < 0.001.

### 5 months of raptor depletion increases fiber size variability but not mean fiber area

Both RAmKO and skeletal muscle fiber-specific mTOR knockout mice (mTORmKO) suffer from a severe, lethal myopathy (Bentzinger et al., 2008; Risson et al., 2009), which is very pronounced at 6 weeks (Risson et al., 2009) and 3 months (Bentzinger et al., 2008) of age, respectively. The RAmKO and mTORmKO myopathy is characterized by smaller fiber sizes, increased fiber size variation, centralized myonuclei, central core-like structures, fibrosis and inflammation. To examine whether iRAmKO mice display similar alterations, we examined cross-sections from TA muscles 21 days, 3 months and 5 months after TAM injection. 21-day iRAmKOs depicted no changes (Supporting Information, Figure S2A), consistent with a previous report (You et al., 2019). After 3 months, despite some fibers from iRAmKO mice appearing more roundish (Figure 3A) and an increased variation in type 2A fiber size distribution and variance coefficient (Briguet et al., 2004) (Supporting Information, Figure S2B and S2 D), overall fiber size, fiber size distribution and variance coefficient were unchanged compared to control littermates (Supporting Information, Figure S2C, S2E, S2F and S2G). Furthermore, we did not detect an increase in the number of central-nucleated fibers (Figure 3C).

**Figure 3.**
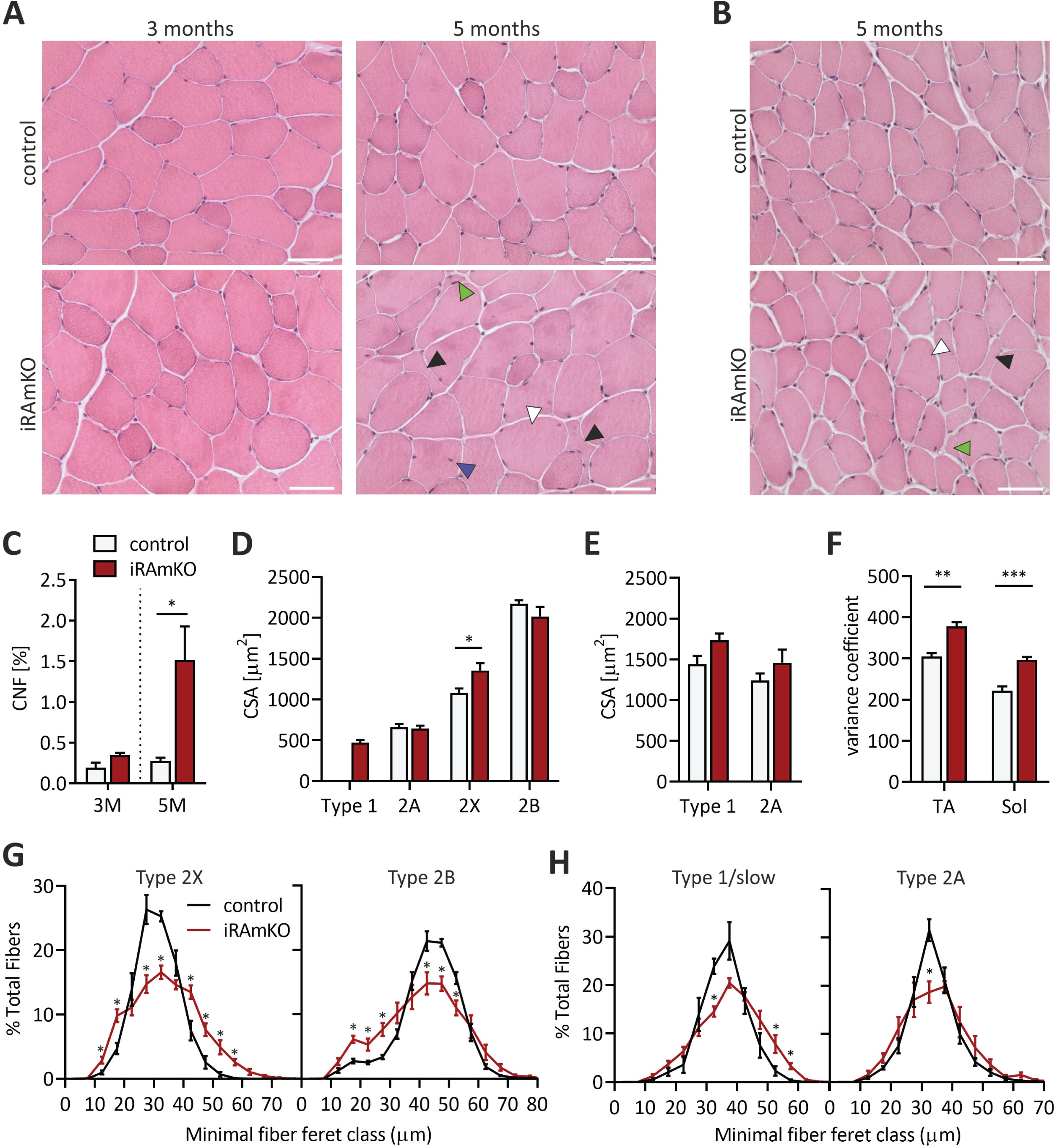
5-months of raptor depletion increases fiber size variability but not mean fiber area. (A) Hematoxylin and eosin (H&E) staining of TA muscle cross-sections from 3- and 5-month iRAmKOs and controls. The black arrows indicate fibers with abnormal morphology, green arrows atrophic fibers, white arrows very large fibers and blue arrows centralized nuclei. (B) H&E staining of *soleus* cross-sections of 5-month iRAmKOs. (C) Quantification of central nucleated fibers (CNF) of the entire TA cross section (using a DAPI and anti-laminin staining). (D) and (E) Average CSA of individual fiber types from the TA (D) and *soleus* (E) of 5-month iRAmKOs. (F) Fiber size distribution as the variance coefficient of 5-month iRAmKOs calculated using the minimal feret’s diameter method (Briguet et al., 2004). (G) Fiber size distribution of type 2X and 2B fibers in the TA of 5-month iRAmKOs and controls. (H) Fiber size distribution of type 1/slow and 2A fibers in the *soleus* of 5-month iRAmKOs and controls. 3-month iRAmKOs n ≥ 3 and 5-month iRAmKOs n = 4. Values represent the mean ± SEM. Significance was assessed using two-tailed unpaired student’s *t*-test: *p < 0.05, **p < 0.01, ***p < 0.001 (for *G* and *H* only *p was used). Scale bars = 50 µm.

The first myopathic features were observable in H&E stainings of the TA muscle from 5-month iRAmKOs. These included higher numbers of very small and large fibers, centrally nucleated fibers and a loss of typical polygonal fiber shape (Figure 3A and 3C). Similar changes were also observed in *soleus* muscle cross-sections from 5-month iRAmKOs (Figure 3B). Mean fiber type-specific and overall fiber area and minimal feret diameter were unchanged in 5-month iRAmKOs in both the TA and *soleus*, with the exception of type 2X fibers in the TA, which were actually hypertrophic (Figure 3D, 3E and Supporting Information, Figure S2H, S2I). On the other hand, there was high numbers of both small and large fibers in TA and *soleus*, as evidenced by a significantly higher variance coefficient (Briguet et al., 2004) (Figure 3F, Supporting Information, Figure S2J, S2K). The fiber size distribution of individual fiber types similarly did not show a clear shift of the mode but presented a lowering of the most abundant fiber size (Figure 3G and 2H; Supporting Information, Figure S2L). This broadening of the fiber size distribution in iRAmKO mice agrees with the increase in the variance coefficient (Figure 3F). In conclusion, raptor depletion in fully-grown muscle does not induce a generalized muscle atrophy, as observed in RAmKOs (Bentzinger et al., 2008) and mTORmKO (Risson et al., 2009), but leads to greater heterogeneity in fiber size. Our data also show that it takes 5 months to detect signs of a myopathy.

### iRAmKO mice have a reduced oxidative capacity and shift the fiber-type composition towards slower fiber types

Another striking phenotype of RAmKO and mTORmKO mice is the significant loss of oxidative capacity, the increase in the number of glycolytic fibers and the change to more slow-type contraction properties (Bentzinger et al., 2008; Risson et al., 2009). In both mouse models, loss of oxidative capacity is based on the downregulation of peroxisome proliferator-activated receptor gamma coactivator 1-α (PGC1α), as transgenic overexpression of PGC1α and pharmacological activation of mitochondrial biogenesis is sufficient to normalize oxidative capacity (Romanino et al., 2011). To assess the metabolic status of iRAmKO mice, we first performed a NADH-tetrazolium (NADH-TR) staining on TA cross-sections. Although mTORC1 signaling was largely abrogated 21 days after TAM injection, the oxidative properties of iRAmKO mice were not different to controls (Supporting Information, Figure S3A). In contrast, 3 months after TAM injection, iRAmKO mice showed a strong decrease in NADH-TR staining (Figure 4A). In line with the NADH-TR staining, gene expression of the transcriptional co-activator PGC1α (*Ppargc1a*) and some of its effector genes, such as myoglobin (*Mb*), cytochrome c (*Cycs*) and cytochrome c oxidase subunit 5B (*Cox5b*), were not altered in 21-day iRAmKOs (Supporting Information, Figure S3B) but were significantly reduced after 3 months (Figure 4B).

**Figure 4.**
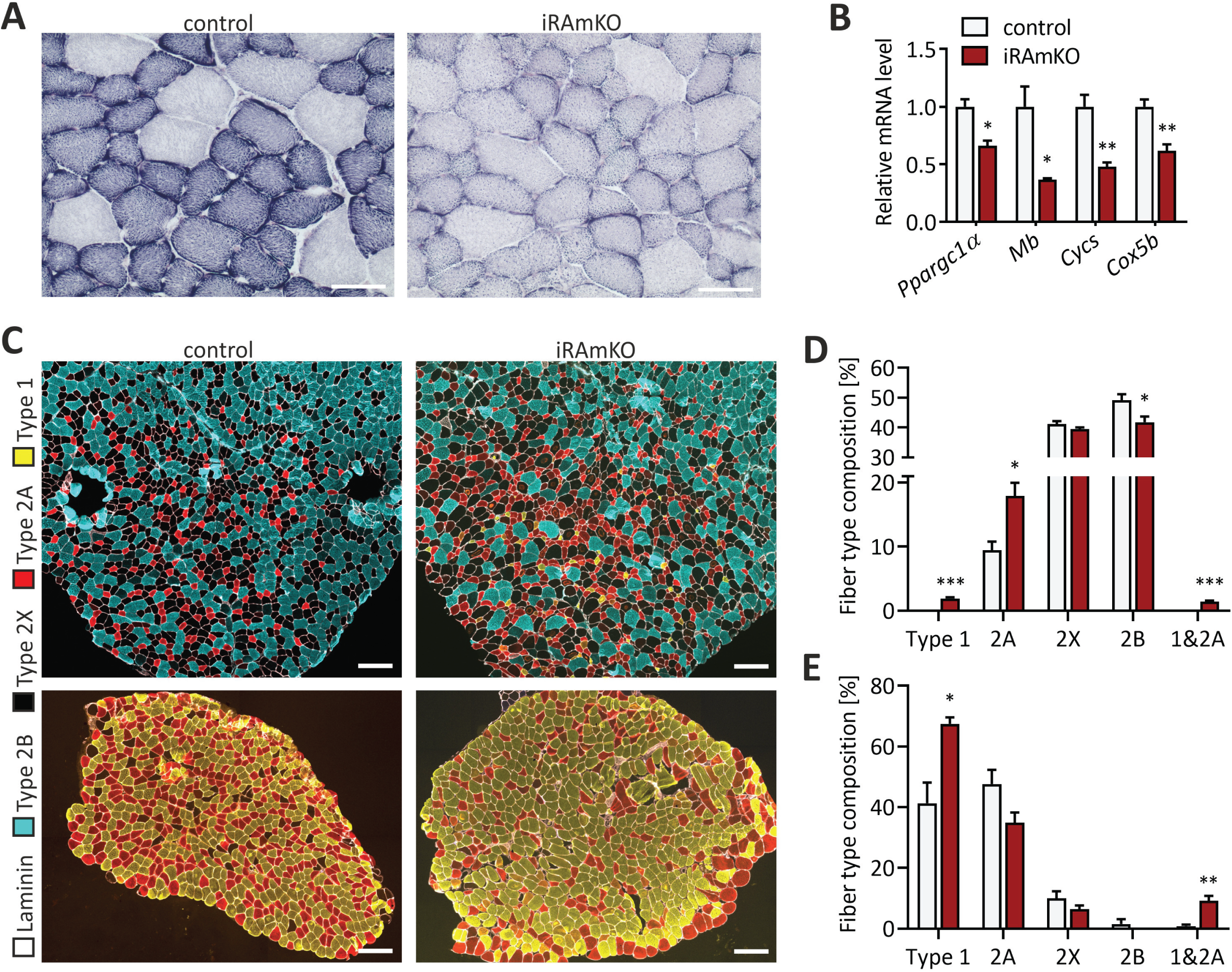
iRAmKO mice have a reduced oxidative capacity and shift the fiber-type composition towards slower fiber types. (A) The activity of oxidative enzymes was examined by NADH-TR staining on TA cross-sections of 3-month iRAmKOs. Scale bar = 50 µm. (B) Relative mRNA levels of *Ppargc1α* (PGC1α), *Mb* (myoglobin), *Cycs* (cytochrome c) and *Cox5b* (cytochrome c oxidase subunit 5B) in the *soleus* muscle of 3-month iRAmKOs. (C) Immunohistochemical (IHC) staining against laminin (white), MyHC type 2B (cyan), MyHC type 2A (red), MyHC type 1/slow (yellow) and type 2X (unstained). The top image is the TA and the lower image is the *soleus* of a 5-month iRAmKO and control. Scale bar = 200 µm. (D) and (E) Quantification of the fiber type composition in TA (D) and *soleus* (E) of 5-month iRAmKOs and their control littermates. n ≥ 3 for 3-month iRAmKOs and n = 4 for 5-month iRAmKOs. Values represent the mean ± SEM. Significance was assessed using two-tailed unpaired student’s *t*-test: *p < 0.05, **p < 0.01, ***p < 0.001.

Fiber-type composition of skeletal muscle is determined by cell intrinsic and extrinsic mechanisms. Muscle fibers formed in the initial wave of myogenesis express largely developmental and slow forms of myosin heavy chains (MyHCs), while muscle fibers formed postnatally express fast MyHCs (Schiaffino et al., 2015). In addition, the pattern of slow and fast MyHCs in the adult is largely determined by motor innervation. Thus, the observed shift in fiber-type composition in RAmKO mice might be due to changes in the intrinsic properties in muscle fibers formed during myogenesis. If this hypothesis were correct, depletion of raptor in adult muscle should not affect fiber-type composition. To test this, we stained muscle cross-sections with antibodies against MyHCs specific for different fiber types. The TA of both 3- and 5-month iRAmKOs presented a two-fold increase of type 2A fibers and a significant reduction of type 2B fibers (Figure 4C and 4D; Supporting Information, Figure S3C). Furthermore, while type 1 fibers are absent in the TA of control mice, ∼2% of fibers in 5-month iRAmKOs are positive for type 1 (Figure 4C and 4D). In line with a fast-to-slow fiber type transition, most type 1 fibers are also positive for 2A fiber type staining (Figure 4D). S*oleus* muscle, which contains ∼40% type 1-positive fibers in control mice, contained more than 60% MyHC 1-positive fibers in 5-month iRAmKO mice (Figure 4C and 4E). As in the TA, *soleus* muscle also exhibited a significant number of type 1 and type 2A-double-positive fibers (Figure 4E). This observation is also confirmed by the mass spectrometry (MS) analysis of the *gastrocnemius* 3 months post-TAM treatment, which showed an increased presence of other proteins and isoforms typically expressed in type 2A and/or slow fiber types, such as Myomesin-3 (Schoenauer et al., 2008), Myosin light chain 6B, Myosin light chain 3 and slow skeletal muscle troponin T (Supporting Information, MS data). Thus, as in RAmKO mice, knockout of raptor in fully-grown muscle causes a reduced oxidative capacity and triggers a switch towards slower fiber types.

### Muscles of iRAmKO mice present slow contractile properties and similar force values to controls

To test how muscle function is affected by long-term mTORC1 inactivation, we performed *ex-vivo* muscle force measurements. The absolute tetanic force of EDL and *soleus* was not significantly reduced in 5-month iRAmKO mice compared to controls (Figure 5A and 5B). Specific force (i.e., force normalized to muscle cross sectional area) in response to stimulation frequency was also very similar in iRAmKO and control mice (Supporting Information, Figure S4A and S4B). Force elicited by a single twitch at 1 Hz was not significantly different in EDL or *soleus* muscle of iRAmKO compared to controls (Figure 5A and 5B). However, the time-to-peak twitch force and half-relaxation time were significantly increased in the *soleus* muscle of iRAmKO mice compared to littermate controls (Figure 5C). In EDL, we could not detect any difference in the time-to-peak but half-relaxation time was significantly increased in iRAmKOs (Figure 5C). Importantly, the EDL muscle of iRAmKO mice was significantly more fatigue-resistant than control EDL (Figure 5D). Our fatigue protocol did not reveal a further increase in fatigue resistance of the iRAmKO *soleus* muscle (Figure 5E). Together, these data show that depletion of raptor in fully grown, adult muscle for 5 months does not lead to a significant reduction in force but does affect muscle contraction properties in line with the fast-to-slow transition in fiber type composition. We thus conclude that the loss of muscle force in RAmKO (Bentzinger et al., 2008) and mTORmKO (Risson et al., 2009) mice is likely a consequence of impaired growth whereas the shift in fiber-type composition may rather be due to mTORC1-driven changes in muscle homeostasis.

**Figure 5.**
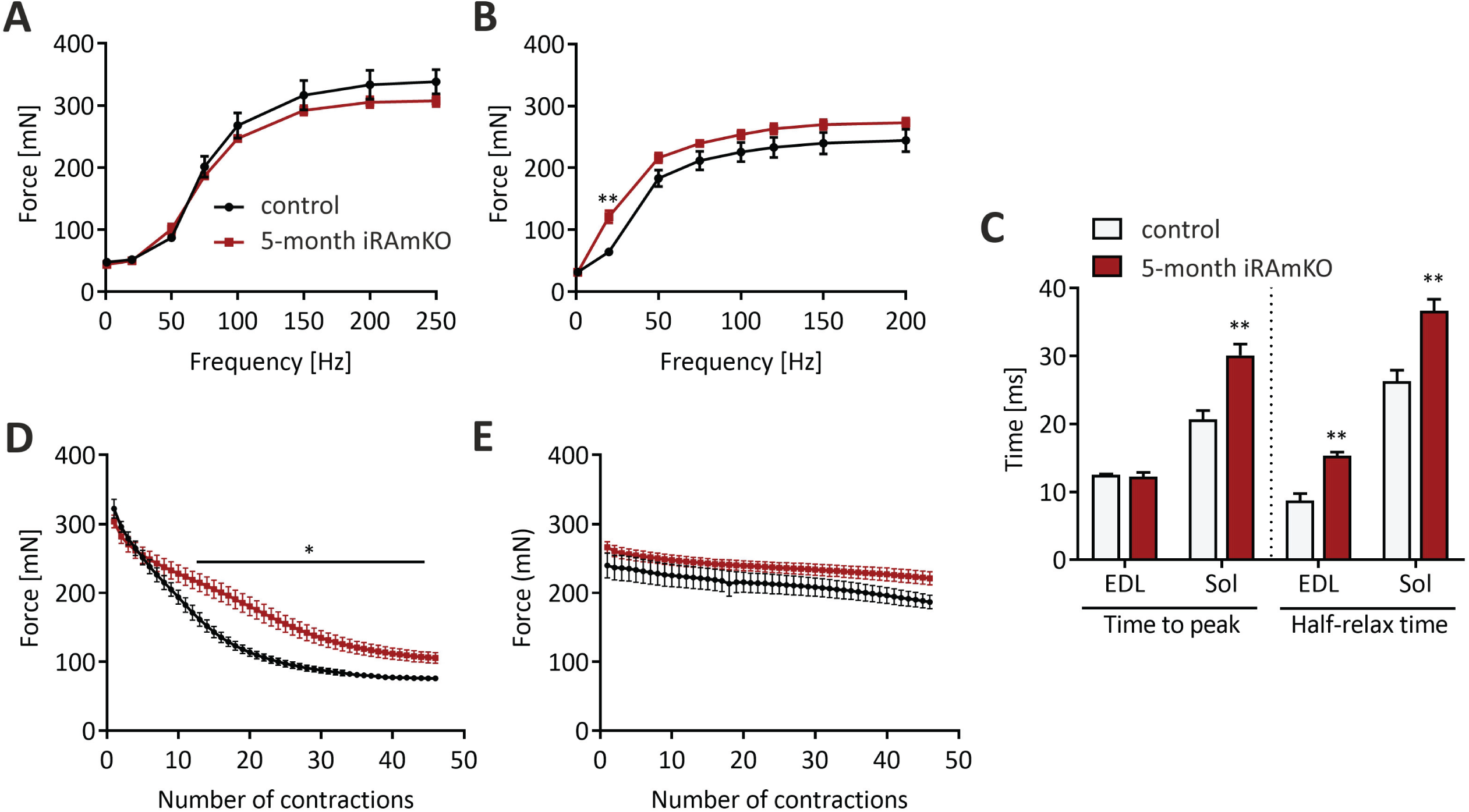
Muscles of iRAmKO mice present slow contractile properties and similar force values to controls. (A) and (B) *Ex vivo* muscle force of the EDL (A) and the *soleus* (B) of 5-month iRAmKO and control mice was measured at different frequencies. (C) Time-to-peak and half-relaxation time of single twitch contractions of EDL and *soleus* muscles in 5-month iRAmKO and control mice. Note the significant prolongation of time components in iRAmKO mice because of the shift to slower fiber types. (D) *Ex vivo* fatigue resistance measured during 6 min with a 200 Hz stimulus every 8 sec. The absolute values are illustrated to show that the muscle force is not decreased. (E) The same fatigue protocol measured in *soleus* muscle at 120 Hz. 5-month iRAmKOs n = 4. Values represent the mean ± SEM. Significance was assessed using two-tailed unpaired student’s *t*-test: *p < 0.05, **p < 0.01, ***p < 0.001

### Removal of mTORC1 leads to a reduction in translational components

mTORC1 controls translation predominantly through phosphorylation of 4E-BP1 and the subsequent release of eIF4E (Thoreen et al., 2012). In iRAmKO mice, inhibition of 4E-BP1, as indicated by its phosphorylation, is strongly reduced (Figure 1C and 1D). We examined whether this also led to greater binding of 4E-BP1 to eIF4E by an m7GTP-pulldown followed by Western blot analysis. The ratio of 4E-BP1 to eIF4E was 2.5-fold higher in iRAmKO mice compared to control, indicating increased inhibition of translation initiation (Figure 6A and 6B). Transcripts that contain a 5’ terminal oligopyrimidine (TOP) or a TOP-like tract show the highest reduction in translation upon 4E-BP1 inhibition (Thoreen et al., 2012). To test whether this was also the case in skeletal muscle, we used mass spectrometry (MS) to quantify the mean level of 40S and 60S ribosomal proteins (RPs), which mostly derive from TOP mRNA (Iadevaia et al., 2008; Thoreen et al., 2012). Indeed, levels of RPs were reduced by ∼24% in mice deficient for raptor (Figure 6C). Similarly, several translation initiation and elongation factors as well as some aminoacyl-tRNA synthetases were strongly reduced (Figure 6D). Together, these data suggest an overall suppression of protein translation in skeletal muscle of iRAmKO mice. To further test this, we performed polysome profiling on the *gastrocnemius* muscle of 3-month iRAmKOs. Typically, a decrease in the translation rate lowers the number of polysomes and increases the number of monosomes and of 40S and 60S ribosomal subunits (Thoreen et al., 2012; Zhang et al., 2019). Indeed, we detected a decrease in the signal from high polysomes in the iRAmKOs (Figure 6E). No increase in the monosome peak nor an increase in 40S and 60S subunits was visible. This lack of an increase of non-polysome ribosomal subunits in iRAmKO muscle is possibly due to the decrease in ribosomal proteins. Hence, removal of raptor does lead to a reduction in global translation, consistent with the results of others (You et al., 2019). Despite this reduction, iRAmKO muscles do not display any atrophy up to 5 months post-raptor depletion (Figure 2D and Figure 3).

**Figure 6.**
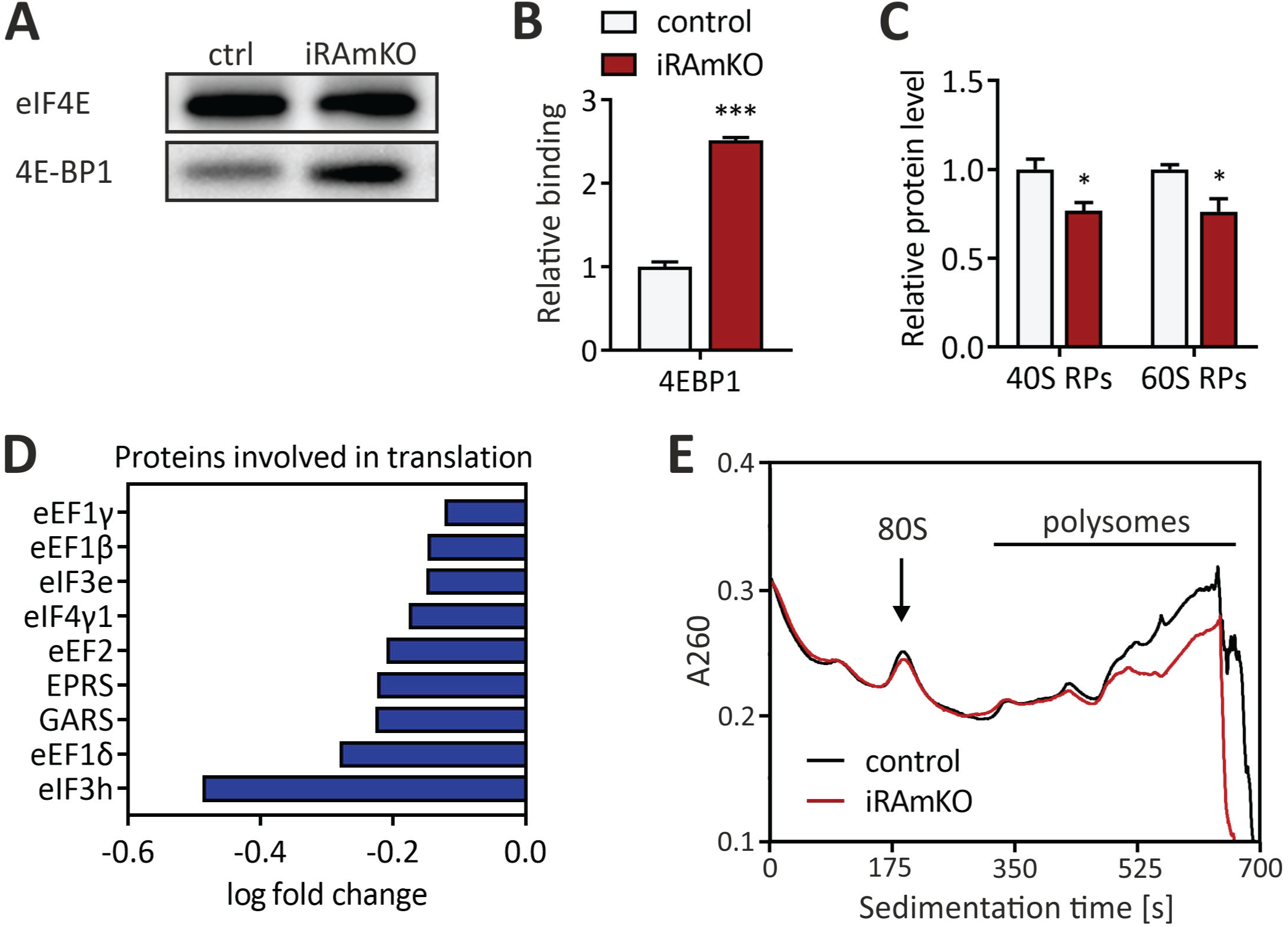
Removal of mTORC1 leads to a reduction in translational components. (A) Western blot of an m7GTP pulldown on *gastrocnemius* lysates of 3-month iRAmKOs with the primary antibodies against eIF4E and 4E-BP1. (B) Quantification of the m7GTP pulldown. The levels of 4E-BP1 were normalised to the amount of eIF4E. (C) Average relative expression levels of the ribosomal proteins obtained from the MS data, separated into 40S and 60S subunits. (D) Significantly decreased proteins involved in translation from the MS data (excluding false discovery rate). (E) Average polysome profiles of *gastrocnemius* from 3-month iRAmKOs and controls. The monosomes are labelled as 80S. 3-month iRAmKOs and controls n = 3. Values represent the mean ± SEM. Significance was assessed using two-tailed unpaired student’s *t*-test: *p < 0.05, **p < 0.01, ***p < 0.001

## Discussion

Using experimental paradigms of hypertrophy, it has been clearly demonstrated that mTORC1 inhibition blunts hypertrophy (Bentzinger et al., 2013; Bodine et al., 2001; You et al., 2019). However, definitive evidence for a role of mTORC1 in the maintenance of adult muscle mass is largely missing. To address this question, we established a mouse model in which we deleted *Rptor* in adult skeletal muscle. Careful characterization of this inducible muscle knockout showed that: (1) all muscle fibers undergo Cre-mediated deletion as indicated by the GFP-positivity, and (2) twenty-one days after tamoxifen injection is sufficient to abrogate mTORC1 signaling. Strong depletion of raptor protein preceded complete inhibition of mTORC1 signaling, as the amount of raptor was already reduced by 70% after 10 days of TAM treatment without any reduction in 4E-BP1 phosphorylation. These data suggest that mTORC1 has quite a long half-live and that low levels of the protein complex are sufficient to maintain normal signaling. Consistent with these data, others have also observed that raptor levels continue to decrease until day 14 after *Rptor* deletion (You et al., 2019). Concomitant with the depletion of raptor, levels of mTOR drop significantly, indicating, as has been suggested from the mTORC1 structure (Aylett et al., 2016), an important role of raptor for the stability/folding of mTOR *in vivo*.

Constitutive, muscle-specific knockout of raptor or mTOR causes a severe, early-onset myopathy that eventually causes the death of the mice. In those mice, body and muscle mass is significantly smaller at the age of 6 to 12 weeks. At the age of 5 months, close to 50% of the RAmKO or mTORmKO mice have died because of the severe myopathy (Bentzinger et al., 2008; Risson et al., 2009). Hence, the most striking result of our current study is the very mild phenotype and the lack of any muscle loss in iRAmKO mice during the 5-month observation period. In 3-month iRAmKO mice, spinal curvature, body and muscle mass were not different from wild-type controls. Furthermore, histological analysis of TA muscle did not show any myopathic signs, such as centralized nuclei, mono-nuclear cell infiltration or fibrosis and overall the variance coefficient was not significantly different. Even after 5 months of raptor depletion, mice still did not show any overt myopathic signs, such as kyphosis, muscle mass loss, or muscle weakness. At this late time point, only early signs of a myopathy started to become visible in muscle histology such as increased fiber size variation and ∼1.5% of central nucleated fibers. These results indicate that the early-onset myopathy observed in RAmKO and mTORmKO mice might be a consequence of impaired muscle hyperplasia or the requirement of juvenile muscle to grow. Moreover, muscle histology shows very mild myopathic signs in 5-month iRAmKO mice that were never as severe as the myopathy described for 3- or 5-month-old RAmKO mice (Bentzinger et al., 2008). We are aware that this experiment does have limitations, in particular because it takes approximately three weeks to eliminate mTORC1 signaling completely. Nevertheless, it provides additional evidence that mTORC1 signaling is not highly critical for skeletal muscle maintenance under sedentary, physiological conditions. Our results are thus different from those obtained when the increase in load causes muscle hypertrophy. In all those paradigms, mTORC1 signaling strongly contributes to hypertrophy (Bentzinger et al., 2013; Bodine et al., 2001; You et al., 2019).

The main role of mTORC1 is to control cell growth by affecting protein synthesis (Ben-Sahra and Manning, 2017). As protein synthesis is important in the S-phase of mitosis, mTORC1 dysfunction strongly affects, although indirectly, fast-dividing cells, including myogenic precursors (Rion et al., 2019). Skeletal muscle is a postmitotic, multi-nucleated tissue, which does not alter cell size fundamentally unless challenged. To assess whether raptor depletion would also affect protein translation, we thus conducted proteomics in *gastrocnemius* muscle from 3-month iRAmKO mice and m7GTP pulldown experiments. Consistent with results obtained in mouse embryonic fibroblasts (Thoreen et al., 2012), ribosomal proteins were strongly and significantly reduced in 3-month iRAmKO mice. The significant loss of ribosomes was accompanied with a decrease in several initiation and elongation factors, such as eIF3, eIF4 and eEF1. Interestingly, all of these downregulated proteins are encoded by TOP mRNAs (Iadevaia et al., 2008). Consistent with a reduction in overall protein synthesis, we observed a relative loss of polysomes in the lysates of iRAmKO muscle. Thus, our results show that TOP mRNAs are also the main target of the translational activity of mTORC1 in skeletal muscle as in other cells (Thoreen et al., 2012; You et al., 2019), including cancer cells (Hsieh et al., 2012). It is thus even more surprising that several months of mTORC1 inactivity does not cause any detectable loss of muscle mass. We hypothesize that a combination of several reasons could explain this lack of phenotype. First, the lowering of TOP mRNA translation may be compensated for by increased translation of other, mTORC1-independent transcripts. Indeed, iRAmKO muscles contain increased levels of several proteins (for example several proteins characteristic of slow muscle fibers; see Supporting Information, MS data). Secondly, the major components of the muscle e.g. myosin and titin, that together constitute ∼34% of muscle mass (Deshmukh et al., 2015), may not be translated cap-dependently and there may not be a great need to synthesize them at high rates because of their low turnover rate under basal conditions (Bates et al., 1983; Bates and Millward, 1983).

While muscle size and myopathic features greatly differed between iRAmKO and RAmKO mice, changes in the metabolism and the fiber-type composition were comparable. A particularly interesting observation is the shift towards slower fiber types. It was previously shown that RAmKOs have a higher proportion of type 1 fibers; however, it was unclear whether this was a consequence of developmental effects (Bentzinger et al., 2008). Myofibers positive for slow MyHC can still be detected up to four weeks after birth in fast muscles, which then vanish later in development (Agbulut et al., 2003; Whalen et al., 1984). Hence, one possibility was that raptor deficiency would prevent this shift to fast type muscle fibers. Our data now show that loss of mTORC1 signaling is sufficient to induce a shift towards slower fiber isoforms in the adult. As raptor is ablated only in muscle, this shift is likely based on muscle-intrinsic, raptor-dependent processes. Interestingly, mTORmKO mice also show this fiber type shift (Risson et al., 2009), indicating that this phenotype is a result of reduced mTORC1 signaling rather than a raptor-specific effect. Two well-described transcription factors involved in the slow fiber switch program are NFATc1 and MEF2 (Kubis et al., 2003; Potthoff et al., 2007; Wu et al., 2000). Levels of MEF2A and MEF2D are both increased in RAmKO muscle (Bentzinger et al., 2008) and NFATc1 has been shown to be negatively regulated by mTORC1 (Zhang et al., 2017). Yet, in both cases, it is unclear whether mTORC1 directly affects these proteins. For example, studies have suggested that the slow removal of calcium from the sarcoplasm characteristic of slow muscle fibers, activates both NFATc1 and MEF2 (Bentzinger et al., 2008). Hence, changes in calcium handling in raptor-depleted muscle may underlie the fiber-type switch. However, the increased removal time of calcium may also be simply a consequence and not a cause of the fast to slow switch. Another protein that may be involved and shows increased activity in raptor-depleted muscle is AKT. Activated AKT inhibits GSK3β, which promotes NFATc1 nuclear entry (Beals et al., 1997). High AKT activity can also lead to increased MEF2 expression (Wiedmann et al., 2005). Thus, constitutively active AKT in iRAmKOs may contribute to the fiber switch. Indeed, time-to-peak and half relaxation time increases in muscles that express a constitutively active form of AKT (Blaauw et al., 2009).

In conclusion, we generated an inducible RAmKO mouse model that allows the analysis of mTORC1 inhibition in fully-grown muscle. We show that raptor knockout in adult muscle has a much weaker effect than expected. Despite a loss of mTORC1 signaling and a significant loss of ribosomes and protein synthesis, muscle mass at no stage becomes significantly less than controls. However, similar to RAmKO and mTORmKO mice, the fast-to-slow fiber-type switch and reduction in oxidative capacity is already visible after 3 months. Thus, this mouse model may allow a better understanding of the mechanisms involved in the anti-aging activity of the allosteric mTORC1 inhibitor rapamycin (Bentzinger et al., 2008; Risson et al., 2009).

## Acknowledgements

We would like to thank the Imaging Core Facility (Biozentrum, Basel-Stadt, Switzerland) and the laboratory of Dr. Christoph Handschin (Biozentrum) that together generated and provided us with the script to analyze fiber-type composition and all parameters of fiber size. This work was financially supported by the Cantons of Basel-Stadt and Basel-Landschaft and by a grant from the Swiss National Science Foundation (#31003A_169845). The authors declare no conflicts of interest.

## Author Contributions

ASH designed and performed experiments, analyzed the data and wrote the manuscript. KC generated the mouse line and helped in the initial phase of the project, LAT performed the polysome profiling, SL performed the muscle force measurements, AS performed the proteomics study, DJH helped analyse the data and discussed the project, MS provided some funding for the project and MAR conceived the project, secured funding, analyzed and discussed data and wrote the manuscript.

## Online supplementary material

Additional supporting information may be found online in the Supporting Information section at the end of the article.

