## Supplemental Figures and Tables for "mTORC1 signaling is not essential for the maintenance of muscle mass and function in adult sedentary mice"

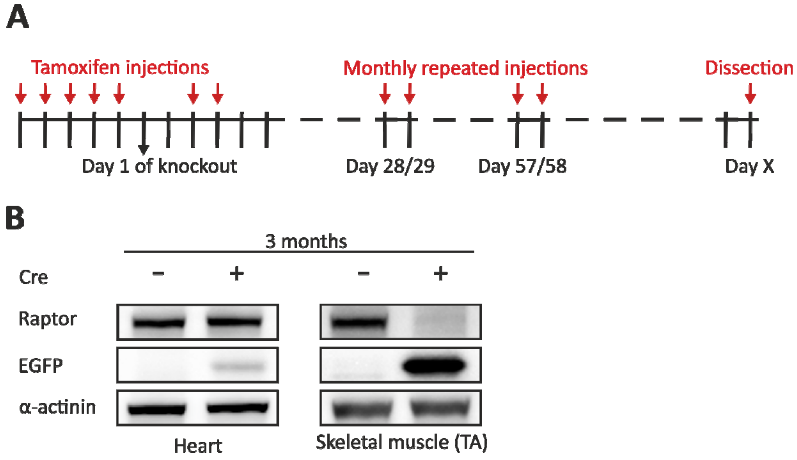


Figure S1. (A) Timeline of TAM injections. (B) Western blot analysis of heart and TA lysates using antibodies directed against the proteins indicated. Note that despite low levels of EGFP expressed in the heart, raptor levels were unchanged, suggesting minimal expression of Cre in this tissue. EGFP was highly expressed in the *tibialis anterior* (TA) muscle and raptor was strongly reduced.

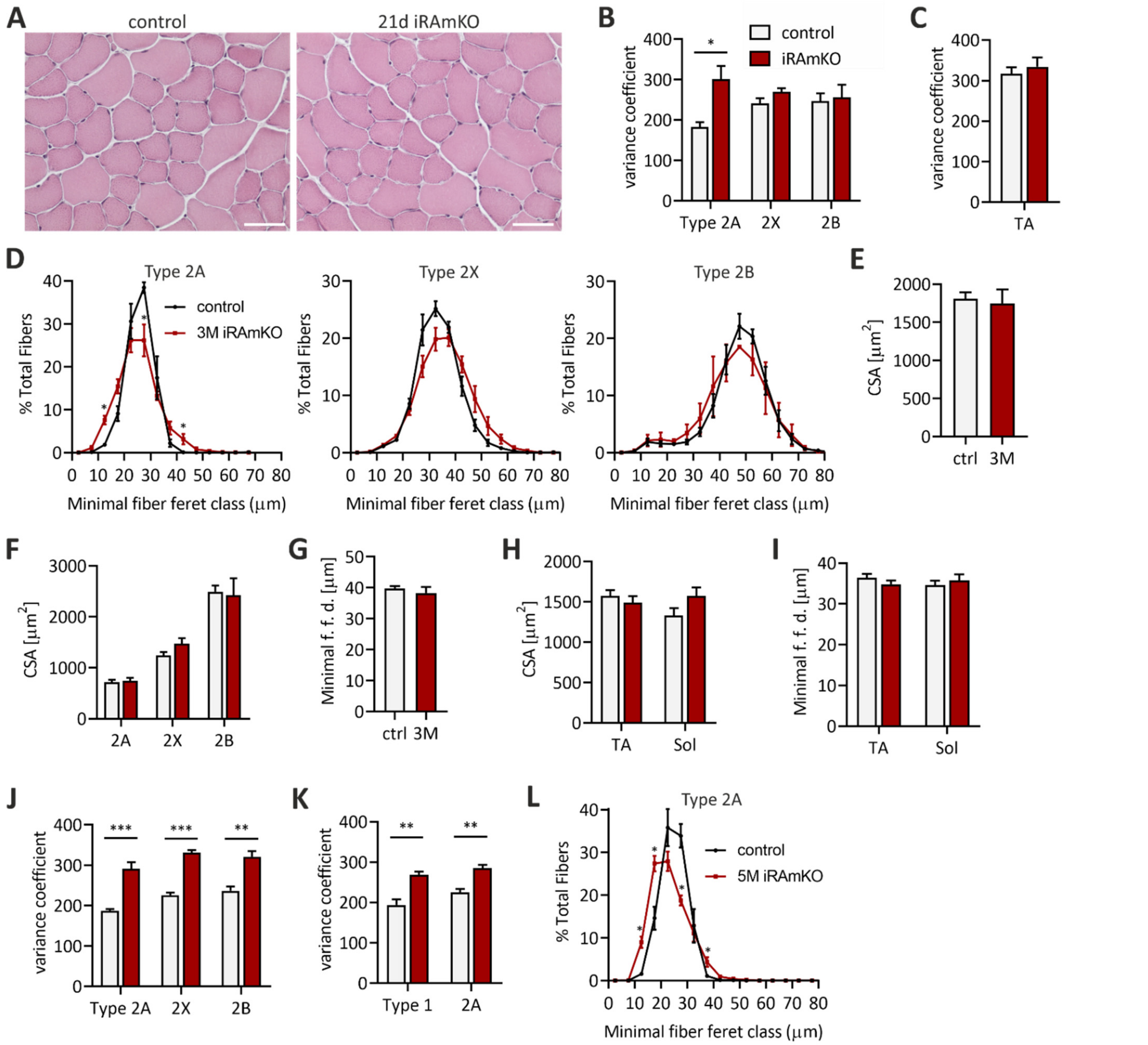


Figure S2. (A) H&E staining of TA cross-sections from 21-day iRAmKOs depicted no changes. Scale bar = 50 μm. (B) The variance coefficient using the minimal feret diameter showed an increase in fiber size distribution for type 2A fibers in the TA of 3-months iRAmKOs. (C) All fiber types of the TA together, however presented no changes in the variance coefficient. (D) Fiber size distribution of type 2A, 2X and 2B fibers from the TA of 3-months iRAmKOs. (E) Average cross-sectional area (CSA) of all fibers and (F) average CSA of individual fiber types from the TA of 3-month iRAmKOs *G*) average minimal fiber feret diameter (f. f. d.) of all fibers from the TA of 3-month iRAmKOs and their control littermates. (H) Average CSA and (I) average minimal f. f. d. of all fibers in the TA and *soleus* (Sol) of 5-month iRAmKOs. (J) Variance coefficient of individual fiber types in the TA and (K) *soleus*. (L) Fiber size distribution of type 2A fibers of the TA from 5-month iRAmKOs. $n \geq 3$ (3M ) and $n = 4$ (5M). Values represent the mean ± SEM. Significance was assessed using two-tailed unpaired student's *t*-test: *$p < 0.05$, **$p < 0.01$, ***$p < 0.001$.

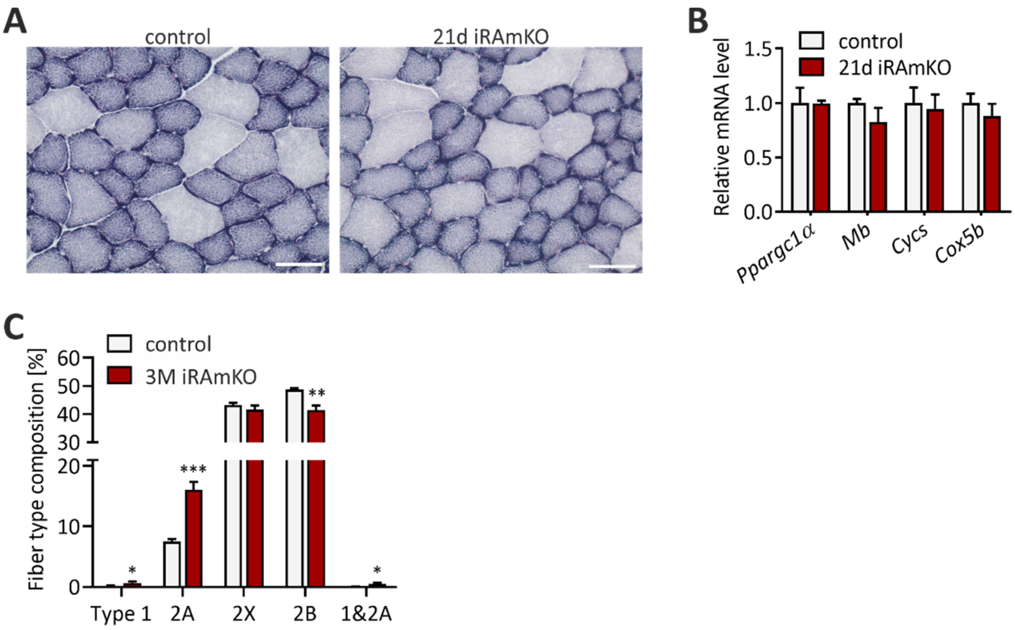


Figure S3. (A) NADH-TR staining of TA cross-sections from 21-day iRAmKOs. (B) Relative mRNA levels of *Ppargc1α* (PGC1α), *Mb* (myoglobin), *Cycs* (cytochrome c) and *Cox5b* (cytochrome c oxidase subunit 5B) in the *soleus* muscle of 21-day iRAmKOs. *C*) Fiber type composition of the TA from 3-month iRAmKOs. n = 4 for 21d iRAmKOs and $n \geq 3$ for 3M iRAmKOs. Values represent the mean ± SEM. Significance was assessed using two-tailed unpaired student's *t*-test: *$p < 0.05$, **$p < 0.01$, ***$p < 0.001$.

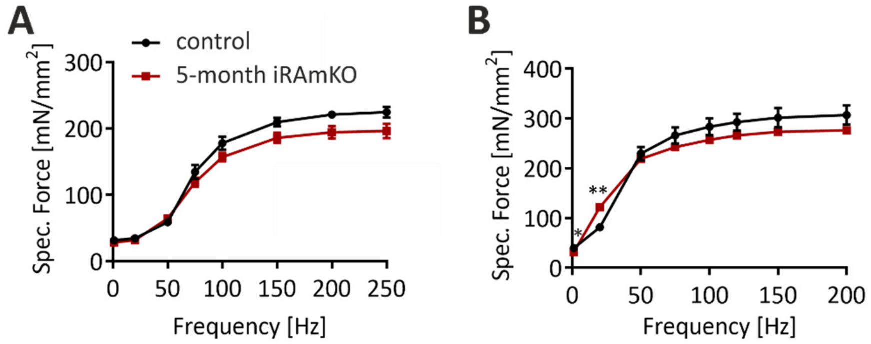


Figure S4. (A) and (B) Specific *ex vivo* muscle force (normalised to the muscle size as explained in material and methods) of the EDL (A) and the *soleus* (B) of 5-month iRAmKOs and controls. 5-month iRAmKOs and controls n = 4. Values represent the mean ± SEM. Significance was assessed using two-tailed unpaired student's *t*-test: *p < 0.05, **p < 0.01, ***p < 0.001.

| Protein | 10 days iRAmKO | | 21 days iRAmKO | | 3 months iRAmKO | |
|---|---|---|---|---|---|---|
| | control | iRAmKO | control | iRAmKO | control | iRAmKO |
| Raptor | 1±0.15 | 0.30±0.02*** | 1±0.13 | 0.11±0.03*** | 1±0.32 | 0.15±0.06** |
| mTOR | 1±0.06 | 0.64±0.02*** | 1±0.14 | 0.39±0.05*** | 1±0.15 | 0.63±0.07* |
| P-mTOR$^{S2448}$ | 1±0.04 | 0.73±0.01*** | 1±0.16 | 0.56±0.04* | 1±0.28 | 0.52±0.09* |
| P-mTOR$^{S2448}$/mTOR | 1±0.09 | 1.13±0.04 | 1±0.05 | 1.38±0.20 | 1±0.16 | 0.86±0.16 |
| 4E-BP1 | 1±0.15 | 1.26±0.20 | 1±0.17 | 1.17±0.29 | 1±0.06 | 1.45±0.13** |
| P-4E-BP1$^{S65}$ | 1±0.24 | 1.73±0.43* | 1±0.13 | 0.64±0.12 | 1±0.12 | 0.16±0.04*** |
| P-4E-BP1$^{S65}$/4E-BP1 | 1±0.25 | 1.37±0.34 | 1±0.17 | 0.51±0.15** | 1±0.12 | 0.11±0.02*** |
| S6 | 1±0.04 | 1.21±0.07** | 1±0.20 | 1.08±0.23 | 1±0.13 | 1.01±0.15 |
| P-S6$^{S240/244}$ | 1±0.14 | 1.15±0.16 | 1±0.42 | 0.87±0.51 | 1±0.15 | 0.71±0.13 |
| P-S6$^{S240/244}$/S6 | 1±0.10 | 0.95±0.13 | 1±0.66 | 0.62±0.25 | 1±0.25 | 0.68±0.02 |
| AKT | 1±0.07 | 1.56±0.06*** | 1±0.20 | 1.05±0.25 | 1±0.25 | 1.42±0.27 |
| P-AKT$^{T308}$ | 1±0.14 | 1.86±0.13*** | 1±0.16 | 3.84±2.68 | 1±0.26 | 7.67±3.11* |
| P-AKT$^{S473}$ | 1±0.12 | 1.83±0.39* | 1±0.14 | 3.37±2.04* | 1±0.32 | 5.19±1.76* |
| P-AKT$^{T308}$/AKT | 1±0.20 | 1.18±0.13 | 1±0.28 | 3.96±3.10 | 1±0.07 | 5.18±1.29** |
| P-AKT$^{S473}$/AKT | 1±0.15 | 1.18±0.30 | 1±0.25 | 2.84±1.08* | 1±0.12 | 3.59±0.63** |

Table S1: All quantifications of the western blots shown in Fig. 1C. All values were normalised on α-actinin and on control samples. 10 and 21-day iRAmKOs n = 4, 3-month iRAmKOs n ≥ 3. Values represent the mean ± SD. Significance was assessed using two-tailed unpaired student's *t*-test: *p < 0.05, **p < 0.01, ***p < 0.001.

| Protein | Heart | | Skeletal muscle (TA) | |
|---|---|---|---|---|
| | control | iRAmKO | control | iRAmKO |
| Raptor | 1±0.24 | 0.95±0.03 | 1±0.32 | 0.15±0.06** |
| GFP | 1±0.28 | 6.00±3.61 | 1±0.49 | 23.15±3.44*** |

Table S2. All quantifications of the western blots shown in Figure S1B. All values were normalised on α-actinin and on control samples. 3-month iRAmKOs n ≥ 3. Values represent the mean ± SD. Significance was assessed using two-tailed unpaired student's t-test: *$p < 0.05$, **$p < 0.01$, ***$p < 0.001$.

**INDIVIDUAL POLYSOME PROFILES**

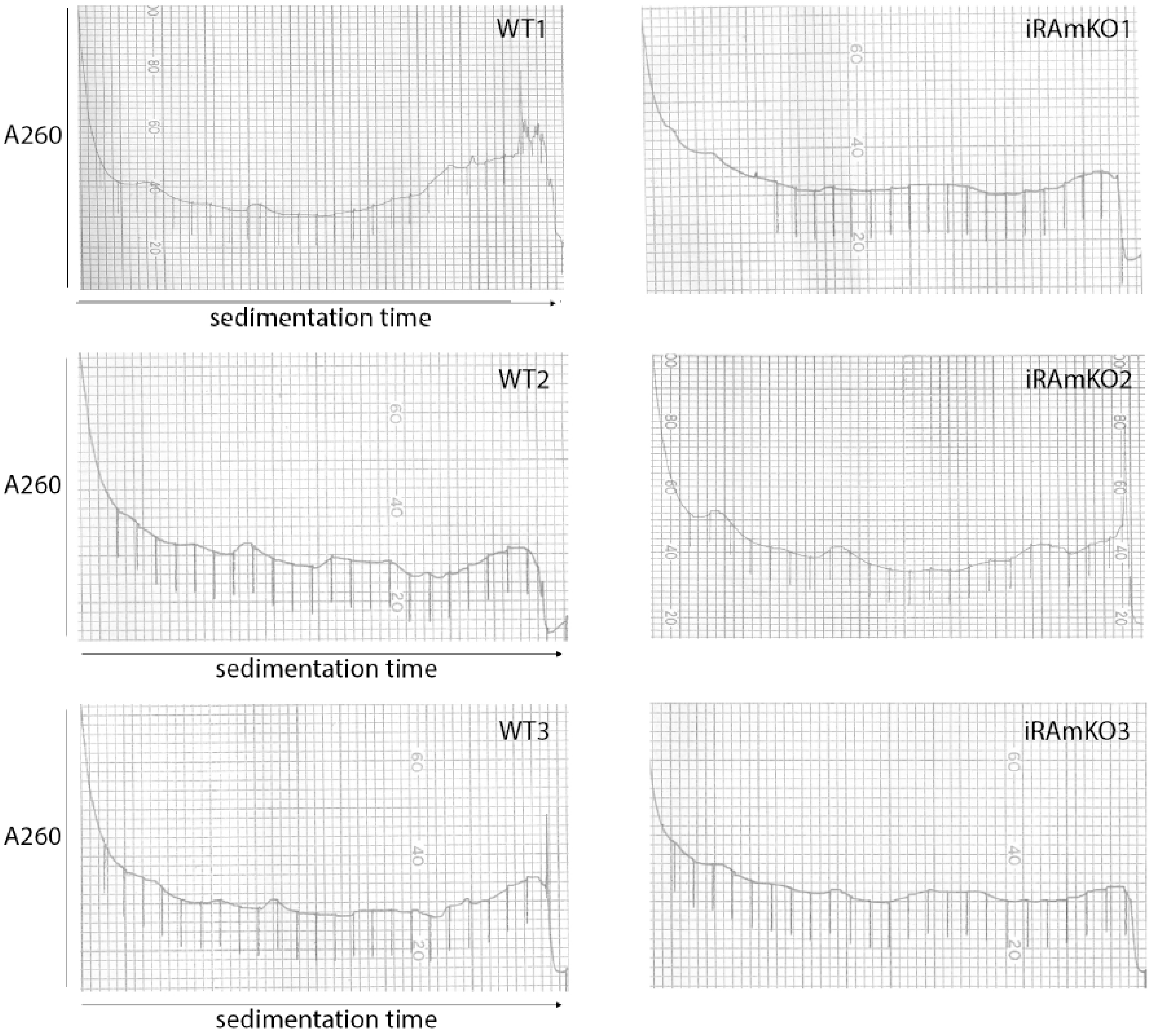
